# β-bursts reveal the trial-to-trial dynamics of movement initiation and cancellation

**DOI:** 10.1101/644682

**Authors:** Jan R. Wessel

**Author notes:** Corresponding Author: Jan R. Wessel, Ph.D., Department of Neurology, University of Iowa Hospitals and Clinics, 444MRC, Iowa City, IA 52242, www.wessellab.org.

## Abstract

The neurophysiological basis of motor processes and their control is of tremendous interest to basic researchers and clinicians alike. Notably, both movement initiation and cancellation are accompanied by prominent field potential changes in the β-frequency band (15-29Hz). In trial-averages, movement initiation is indexed by β-band desynchronization over sensorimotor sites, while movement cancellation is signified by β-power increases over (pre)frontal areas. However, averaging misrepresents the true nature of the β-signal. As recent work has highlighted, raw β-band activity is characterized by short-lasting, burst-like events, rather than by steady modulations. To investigate how such β-bursts relate to movement initiation and cancellation in humans, we investigated scalp-recorded β-band activity in 234 healthy subjects performing the Stop-signal task. Four observations were made: First, both movement initiation and cancellation were indexed by systematic, localized changes in β-bursting. While β-bursting at bilateral sensorimotor sites steadily declined during movement initiation, β-bursting increased at fronto-central sites when Stop-signals instructed movement cancellation. Second, the amount of fronto-central β-bursting clearly distinguished successful from unsuccessful movement cancellation. Third, the emergence of fronto-central β-bursting coincided with the latency of the movement cancellation process, indexed by Stop-signal reaction time. Fourth, individual fronto-central β-bursts during movement cancellation were followed by a low-latency re-instantiation of bilateral sensorimotor β-bursting. These findings suggest that β-bursting is a fundamental signature of the motor system, reflecting a steady inhibition of motor cortex that is suppressed during movement initiation, and can be rapidly re-instantiated by frontal areas when movements have to be rapidly cancelled.

**Significance Statement:** Movement-related β-frequency (15-29Hz) changes are among the most prominent features of neural recordings across species, scales, and methods. However, standard averaging-based methods obscure the true dynamics of β-band activity, which is dominated by short-lived, burst-like events. Here, we demonstrate that both movement-initiation and cancellation in humans are characterized by unique trial-to-trial patterns of β-bursting. Movement initiation is characterized by steady reductions of β-bursting over bilateral sensorimotor sites. In contrast, during rapid movement cancellation, β–bursts first emerge over fronto-central sites typically associated with motor control, after which sensorimotor β–bursting re-initiates. These findings suggest a fundamentally novel, non-invasive measure of the neural interaction underlying movement-initiation and –cancellation, opening new avenues for the study of motor control in health and disease.

## Introduction

Activity in the β-frequency band (15-29Hz) is a prominent constituent of the neural field potential. It can be observed at spatial scales ranging from extracellular to scalp recordings, in species ranging from rodents to humans, and using methods ranging from intracranial recordings to magnetoencephalography [1-4]. The β-frequency plays a particularly important role in the functioning of the motor system. In particular, during movement initiation, a prominent desynchronization of β-band activity is clearly observable over sensorimotor areas [5, 6]. In contrast, the rapid cancellation of movement is accompanied by β-power increases over (pre-)frontal cortical areas generally implicated in cognitive control [7-11]. Moreover, movement-related changes in β-power can also be observed in extrapyramidal parts of the motor system, including the basal ganglia [12-15], where abnormal β-rhythms are prominently observed in movement disorders such as Parkinson’s Disease [16-19].

Recent studies of raw, unaveraged β-band activity, however, have led to a significant reappraisal of the nature of this β-band activity. While trial-averaging approaches suggest that movement-related changes in the β-band reflect steady (de)synchronizations that stretch over several hundred milliseconds, unaveraged β-band activity is primarily characterized by rapid burst-like events, which typically last less than ∼150ms [20-22]. While these burst-events appear as slow-evolving (de)synchronizations in the trial-average, analyses of single-trial data have found that the simple presence or absence of these β-bursts, rather than overall changes in β-power, is the most reliable predictor of trial-to-trial behavior [3].

Here, we therefore investigate the characteristics of single-trial β-bursting in humans during both the initiation, and particularly, during the rapid cancellation of movement. A large sample of healthy human participants (N=234) performed the stop-signal task [23, 24], a motor task that includes both instances of movement initiation (following a Go-signal) and movement cancellation (on trials that include a subsequent Stop-signal). We used non-invasive scalp-EEG recordings to investigate how β-bursting on individual trials indexes both processes and the interaction between them. Specifically, we investigated five questions: 1. Is human β-band activity during movement *burst-like*? 2. Are there systematic *topographical* patterns of β-bursts during both movement initiation and –cancellation? 3. Are movement initiation and cancellation accompanied by systematic *temporal* patterns of β-burst activity? 4. Do specific patterns of β-burst activity distinguish *successful from unsuccessful* movement cancellation? 5. Are there systematic *relationships* between initiation and cancellation-related changes in β-bursting when movements have to be rapidly stopped?

## Results

### Systematic spatiotemporal patterns of β-bursts characterize both movement initiation and cancellation

Figure 1a shows that after Go-signals (which prompt the start of movement initiation), bilateral sensorimotor sites (electrodes C3 and C4) initially show localized β-bursting, which immediately begins to decrease as time gets closer to movement execution, resulting in a significant linear trend (Figure 1b, d, linear trend for left-hand responses at C4: Z = -3.63, p = .00028, left-hand responses at C3: Z = -3.22, p = .001, right-hand responses at C4: Z = -2.81, p = .005, right-hand responses at C3: Z = -3.23, p = .0013). Furthermore, in the lead-up to response execution, this pattern lateralizes, with sites contralateral to the response hand showing a stronger sustained reduction in β-bursting (significant lateralization at p < .0001 (FDR-corrected) at time five consecutive windows from 325 to 575ms for electrode C4, t(233) = -4.91, p = 1.75*10^-06^, d = .38, t(233) = -8.25, p = 1.2*10^-14^, d = .63, t(233) = -6.74, p = 1.24*10^-10^, d = .52, t(233) = -6.83, p = 7.37*10^-11^, d = .61, t(233) = -5.75, p = 2.81*10^-08^, d = .5; and at three consecutive time windows from 325 to 475ms for electrode C3, t(233) = 5.28, p = 2.92*10^-07^, d = .41, t(233) = 6.02, p = 6.6*10^-09^, d = .49, t(233) = 4.81, p = 2.65*10^-06^, d = .37). An inspection of individual trial data shows that the single-trial β-band signal is indeed dominated by short, burst-like events, rather than by steady modulations (Figure 1c, e; plots of single trial data for each individual participant can be found in the Supplementary Materials).

**Figure 1.**
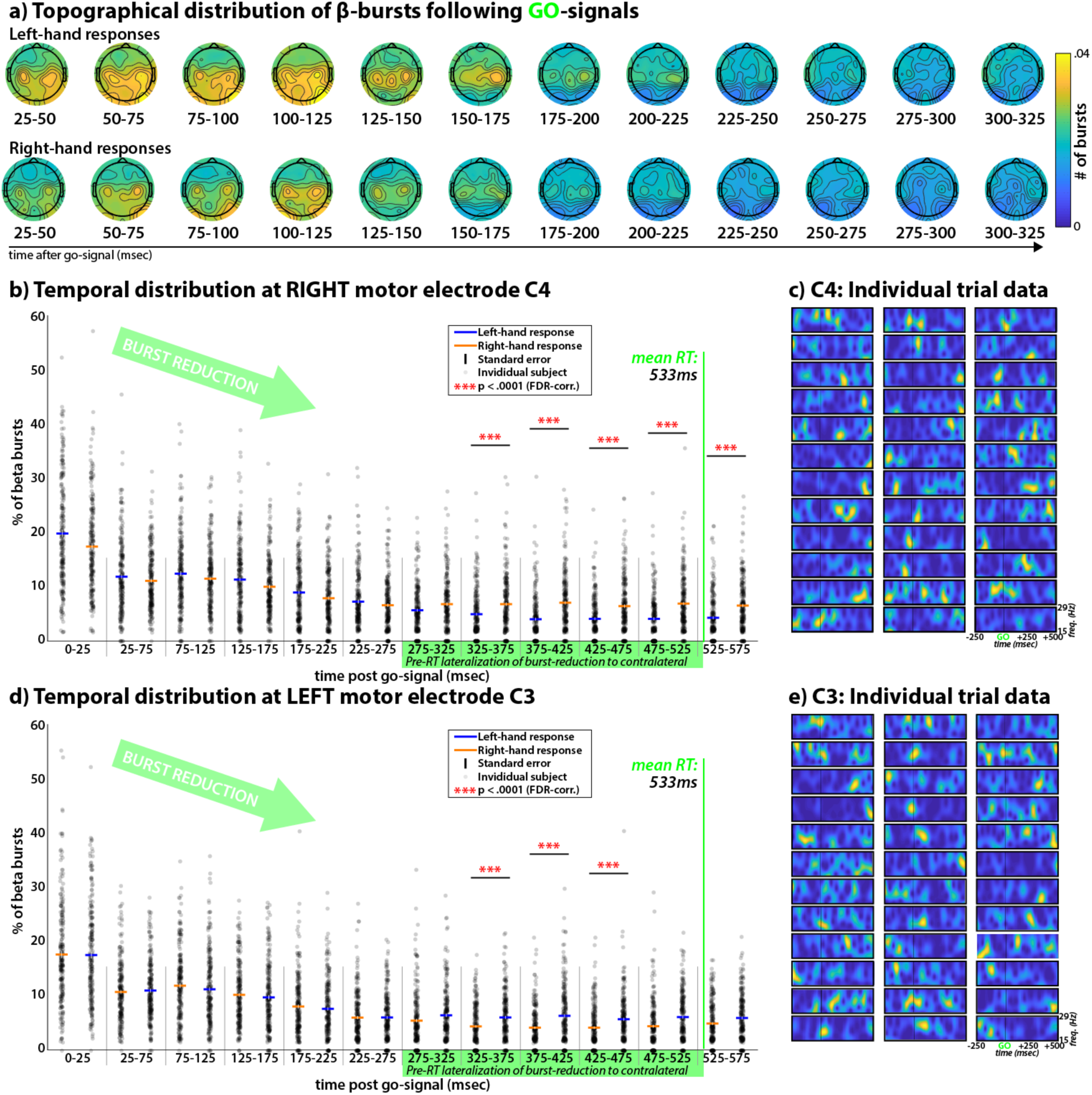
β-burst properties during movement initiation on Go-trials. A) Topographical distribution of the number of β-bursts on the scalp at different consecutive time-windows following the Go-signal, for both left-hand (top) and right-hand (bottom) responses. There are visible bilateral peaks over electrode sites C3 and C4 in both conditions until about 200ms following the Go-signal. B) Temporal distribution of β-burst numbers following the Go-Signal at right lateral electrode C4 during both left- and right hand responses. A significant linear trend can be observed, such that the number of β-bursts steadily decreases following Go-signal onset. Moreover, there is a significant lateralization of this effect starting at 325ms following the Go-signal, such that the β-burst numbers for the contralateral hand keep diminishing while the number asymptotes earlier for the ipsilateral response hand. C) Individual trial β-band data at electrode C4 from one representative subject, clearly showing burst-like β-events. D) As B, but for left-lateral electrode C3. E) As C, but for C3.

Figure 2a shows that in the time period following Stop-signals (which followed Go-signals on 1/3 of all trials at a variable delay and prompted the participants to attempt to stop their pending movement), no coherent spatiotemporal organization of the rate of β-bursts can be observed until around 200ms after the stop-signal. At that point, a clear radial fronto-central topographical distribution emerges, centered around electrode FCz. Just like Go-signal-related activity at C3 and C4, single-trial Stop-signal-related activity at FCz clearly shows the presence of β-bursting (Figure 1d).

**Figure 2.**
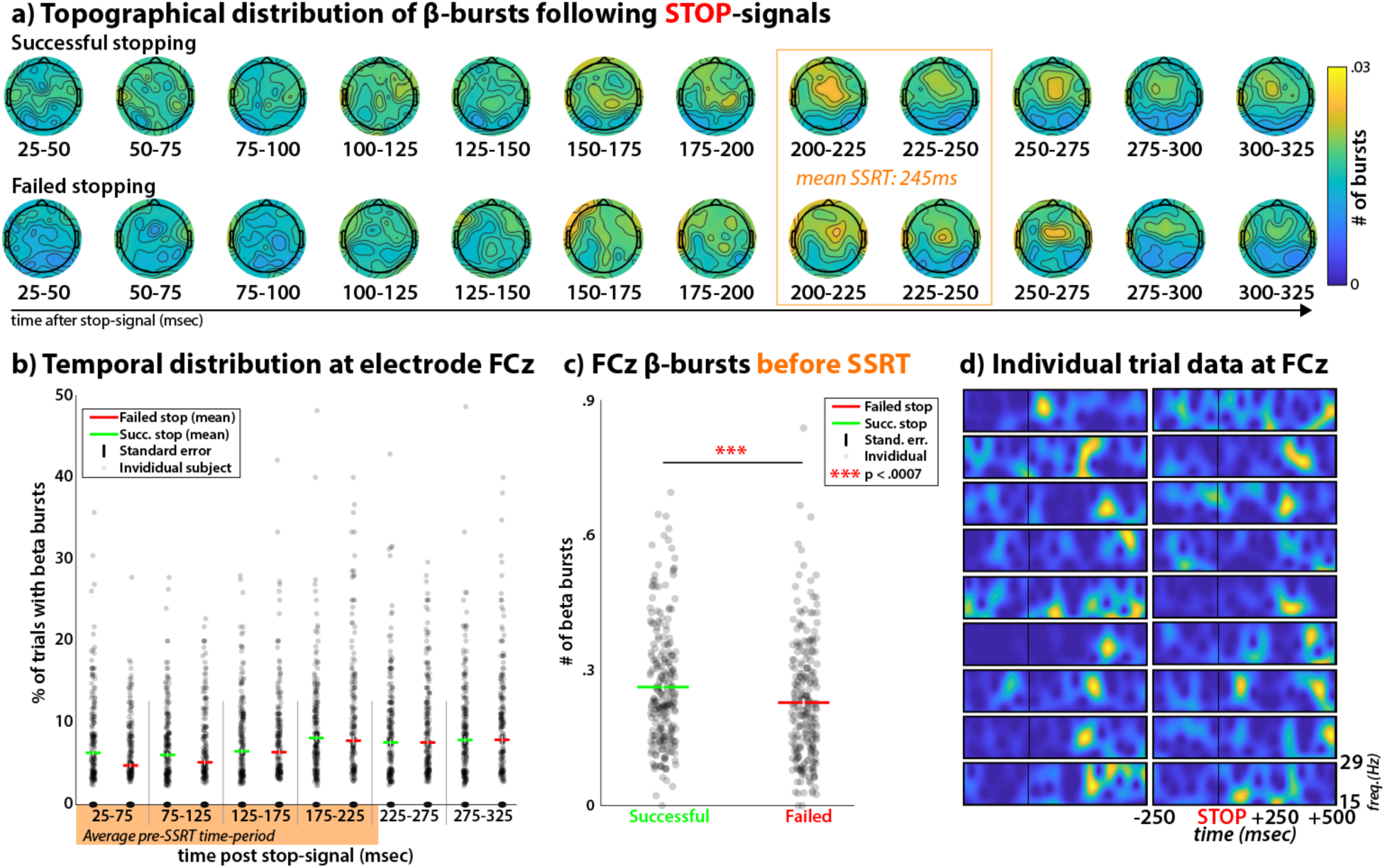
β-burst properties during movement cancellation on Stop-trials. A) Topographical distribution of the number of β-bursts on the scalp at different consecutive time-windows following the Stop-signal, separately for successful (top) and failed (bottom) Stop-trials. In the time-window towards the end of SSRT (orange window), a clear fronto-central organization of β-bursting centered around electrode FCz is observable. B) Temporal distribution of β-burst numbers following the Stop-Signal at fronto-central electrode FCz. C) Comparison of the number of β-bursts between successful and failed Stop-trials in the Stop-signal-to-SSRT period for each subject. Successful Stop-trials are accompanied by a significantly larger number of β-bursts. D) Individual trial β-band data at electrode FCz from one representative subject, clearly showing burst-like β-events.

### Fronto-central β-bursting indexes successful movement cancellation

Behavior in the Stop-signal task was typical (mean Go-trial reaction time: 533.51ms, SEM: 6.6; failed Stop-trial reaction time: 459.59ms, SEM: 5.79; stop accuracy: .52, SEM: .002, Stop-signal delay: 282.42, SEM: 7.91; Stop-Signal reaction time: 244.98ms, SEM: 3.62).

The increase of fronto-central β-bursting after Stop-signals (compared to Go-signals) is highly significant (Figure 3 shows the topographical difference plots between the distribution of β-bursts following Stop- vs. Go-trials, thresholded at p < .0001, FDR-corrected). Moreover, the time after the Stop-signal at which this fronto-central organization of β-bursting develops overlaps with the end of SSRT (∼245ms). The period before the end of SSRT is the exact time period during which neural activity reflecting movement cancellation should be maximal according to computational models [25]. Moreover, a direct comparison of the pre-SSRT time period in each individual participant (i.e., the time range between the Stop-signal and the end of that participants’ SSRT) revealed that successful Stop-trials yielded an increased rate of β-bursts at FCz compared to failed Stop-trials (t(233) = 3.47, p < .0007, d = .26, Figure 2b, c).

**Figure 3.**
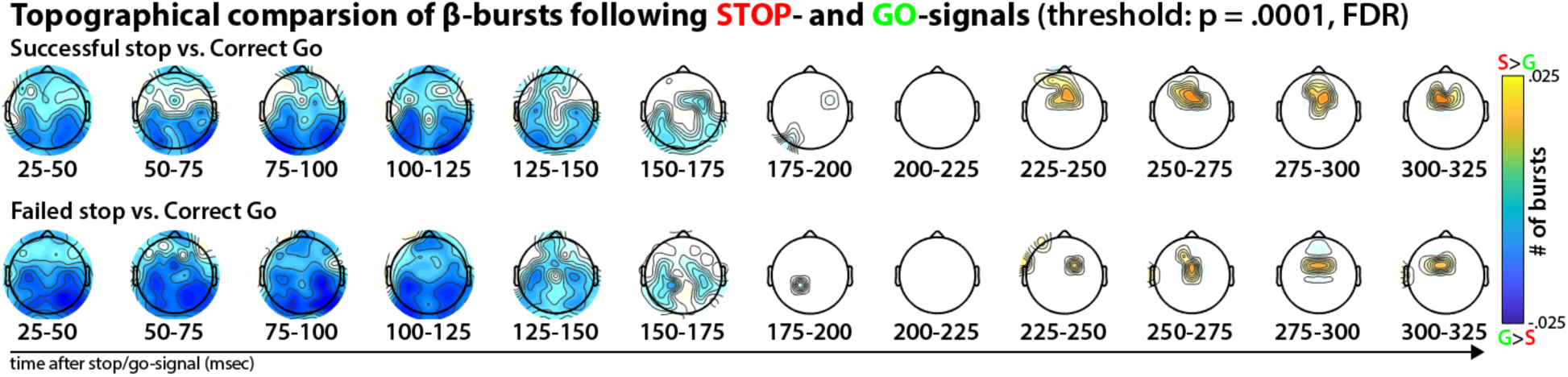
Statistical comparison of β-burst topographies following Stop- and Go-signals, separately for successful (top) and failed (bottom) Stop-trials. Thresholded for significance at p < .0001, FDR-corrected for multiple comparisons. It is evident that while movement initiation (Go-trials) is accompanied by a significantly increased β-burst count at lateral electrodes until ∼175ms after the Go-signal, movement cancellation is accompanied by a significantly increased β-burst count at fronto-central electrodes, starting at ∼225ms after the Stop-signal.

### Fronto-central β-bursts are followed by increased bilateral sensorimotor β-bursting

Figure 4a shows the temporal development of β-bursting at sensorimotor sites ipsi- and contralateral to the to-be-stopped movement on successful stop-trials. Importantly, these plots are time-locked to the latency of the first fronto-central β-burst event that occurred within the pre-SSRT time period. This is the time period during which the Stop-process should be active according the race-models of the Stop-signal task [23, 25, 26]. These plots show a significant increase in bilateral sensorimotor β-bursting within 25ms of the first fronto-central β-burst. To evaluate significance, these values were compared to matched time-periods on successful stop-trials without fronto-central β-bursts in the pre-SSRT period (exact values for the pairwise comparisons between trials with and without β-bursts: Z = 7.81, p = 5.65*10^-15^ and Z = 4.36, p = 1.28*10^-05^ for the two significant time-windows for contralateral sites and Z = 8.06, p = 7.7*10^-16^ and Z = 4.8, p = 1.55*10^-06^ for ipsilateral sites). A topographical representation of this effect reveals that the increase in β-bursting following the first fronto-central β-burst is localized to bilateral sensorimotor sites (in addition to the fronto-central electrodes surrounding FCz, Figure 4b).

**Figure 4.**
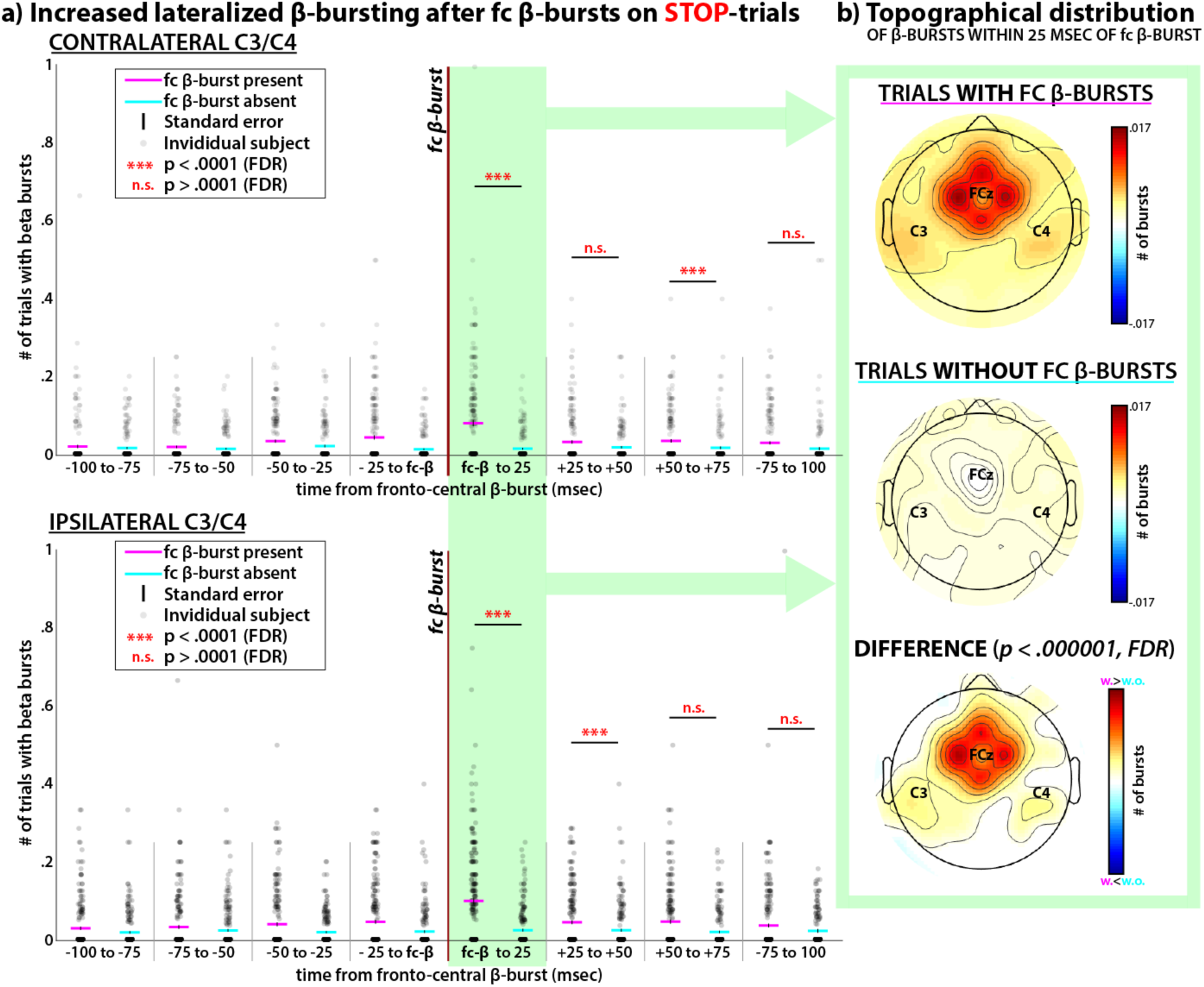
Interaction between fronto-central and lateralized β-bursting during successful movement cancellation. A) Number of contra- and ipsilateral β-bursts at electrodes C3/4 surrounding fronto-central β-bursts during the Stop-signal-to-SSRT period. During the time window 25ms following the fronto-central β-burst (green highlighting), both contra- and ipsilateral β-bursting over electrodes C3/C4 was significantly increased. B) Topographical representation of β-bursting in the highlighted time period, 25ms following the fronto-central β-burst event, showing that increases in β-burst activity are localized to bilateral sensorimotor sites (in addition to fronto-central sites surrounding FCz).

## Discussion

The current set of findings shows that β-bursting on individual trials is tightly related to movement in humans. Prior to movement initiation, bilateral sensorimotor sites showed localized patterns of β-bursting, which may represent an inhibited state of the motor system [4, 11, 27]. This β-bursting is then steadily reduced when a Go-signal is presented and a movement is initiated (reflecting a net-disinhibition of the motor system). Notably, this reduction in burst-rates lateralized just prior to movement execution, which mirrors prior observations of lateralized movement-related β-desynchronizations in the trial average [6, 28, 29], and likely explains these patterns.

Subsequently, when inhibition had to be rapidly reinstated – i.e., when already initiated movements had to be suddenly cancelled following Stop-signals – β-bursting significantly increased at fronto-central scalp sites. This fronto-central increase in β-bursting showed its most coherent spatiotemporal organization at fronto-central sites during time period towards the end of SSRT. This is the exact time period that, according to a popular computational model of the Stop-signal task, should yield the highest amount of inhibitory activity [25]. Finally, one highly notable pattern in our data was that individual instances of fronto-central β-bursting were followed by a rapid (<25ms) re-instantiation of β-bursting over sensorimotor sites, both ipsi- and contralateral to the to-be-stopped movement. We propose that this reflects a low-latency re-instantiation of inhibition at the level of the motor system, triggered by a fronto-central control signal (reflected in the fronto-central β-burst events).

These features of our data show intriguing overlap with several of the proposed properties of the neural cascade underlying movement cancellation that was identified in prior work using measures other than β-bursting. First, the topographical distribution and temporal evolution of the fronto-central β-burst reported here parallels the properties of the Stop-signal P3, an event-related potential that has been proposed to reflect motor inhibition in trial-averaging studies of phase-locked event-related EEG activity [30-32]. Second, trial-averaged power in the β-band has been repeatedly implicated in movement cancellation in studies of trial-averaged time-frequency activity [7-10, 13]. This includes intracranial recordings from sites that could well underlie the fronto-centrally distributed pattern of β-bursting observed in our study, including the pre-supplementary motor area [33]. Previous studies may have missed these β -burst patterns due to trial-averaging. Third, the prominent race-model of the Stop-signal task proposes that behavior in this task can be modeled by a race between two processes; one working towards the execution of the motor response and triggered by the Go-signal (the Go-process), and one working towards the cancellation of that response and triggered by the Stop-signal (the Stop-process [23]). Some controversy still remains regarding whether these two processes operate independently, or whether the Stop-process, once initiated, directly influences the Go-process [25, 26, 34, 35]. Our results show that stopping-related activity at fronto-central sites is immediately followed by a re-instantiation of β-bursting over sensorimotor sites – the same signature whose initial reduction represents the start of the Go-process (Figure 4). This indicates that the activity of the Stop-process is followed by a substantial change in the neural representation of the Go-process, thus speaking in favor of the interactive race-model of movement cancellation. Finally, the pattern of β-bursts during movement cancellation in our study dovetails well with a fourth aspect of the existing literature. Notably, the Stop-related increase in sensorimotor β-bursting following fronto-central β-bursts was equally observable at both ipsi- and contralateral sensorimotor sites. This parallels existing findings from the motor physiology literature on the Stop-signal task, which have used motor evoked potentials (a signature of cortico-spinal excitability of specific motor tracts) to show that rapid movement cancellation is non-selective. Specifically, when movements are rapidly cancelled, redutions of cortico-spinal excitability are not limited to the specific motor effectors that have to be stopped. Instead, suppression of the entire motor system is observed, even at task-unrelated muscles [36-39]. The simultaneous, rapid re-activation of β-bursting over *both* sensorimotor cortices after fronto-central Stop-related β-bursts could be the neurophysiological expression of that same non-selective property in our current dataset.

The current study has several key implications for future research, and motivates several immediate follow-up experiments. First, the precise neural origin of movement-related β-bursts needs to be investigated. Computational modeling has suggested that these events may results from the integration of near-synchronous bursts of excitatory synaptic drive, targeting the dendrites of pyramidal neurons in specific cortical layers [22]. Whether this physiological property dovetails with the features of specific brain regions that are the purported neural generators of the scalp-recorded data reported here remains to be tested. Second, if β-bursting over sensorimotor sites is indeed indicative of a ‘tonic’ inhibitory mode of the motor system – one that is reduced during movement initiation and re-initiated during movement cancellation – it would make it an interesting target for investigations of tonic, preparatory control activity, e.g., that found in proactive inhibition tasks [40-45]. For example, increases in proactive inhibition in task contexts with higher relative likelihoods of Stop-signals could result in increased bilateral sensorimotor β-bursting. Third, there is great interest in pathological features of the β-frequency band in movement disorders, especially Parkinson’s disease [16-19]. To date, most studies of β-band activity in PD are still based on trial-averages. However, recent studies have already used trial-level β-burst measurements in subcortical areas of the basal ganglia to identify gait problems in PD [46] and have investigated the effect of deep-brain stimulation on subcortical β-burst [47]. Future studies may aim to investigate the relationship between abnormal patterns of β-bursting on the scalp and the specific impairments in movement cancellation that are commonly observed in PD [48-50]. Fourth, β-bursts may provide a new window into the interactions between the subcortical aspects of the extrapyramidal motor system and the cortical aspects underlying higher levels of movement-planning and –control. Several studies have already described β-bursting in several basal ganglia nuclei [17, 20, 21, 51]. It is tempting to assume a correspondence between β-bursting in these brain regions and the cortical regions that are likely underlying the patterns observed in the current study. Indeed, β-bursting may be a ‘universal’ language of the motor system, signifying distributed processing throughout the both pyramidal and extrapyramidal motor pathways. Finally, investigating fronto-central β-bursts could provide a fruitful test-bed for studies of motor inhibition across many different psychological paradigms. As has been proposed elsewhere, inhibitory control may be a rather universal control mechanism, which could be involved in regulating behavior after unexpected events [52-54], response-conflict [55, 56], error processing [57, 58], and many other control processes. The current study suggests that even already-existing datasets of experimental tasks that operationalize these processes and measure the neurophysiological indices may be worth re-investigating with a focus on β-band bursting.

Given its nature as the first investigation of single-trial β-bursting during human movement cancellation (and –initiation), the current study cannot answer all potential questions, and has several shortcomings. First, since the neural activity in the current study was non-invasively recorded from the scalp, only very limited conclusions can be made regarding the exact origins of the observed β-bursts. Second, while the fronto-central β-bursts observed in the current study are clearly related to the success of movement cancellation, they are also clearly neither necessary nor sufficient. In fact, Figure 2c shows that even failed Stop-trials include fronto-central β-bursts (though at a reduced rate). Furthermore, the figure also shows that the majority of successful Stop-trials did not include a β-burst event (at least in the pre-SSRT period). One possible reason for this is that typically, not all Stop-trials in the Stop-signal task actually involve the initiation of a Go-response, which is a requirement for motor inhibition [44, 59, 60]. This is especially true on trials on which the Stop-signal delay is very short, giving little time for significant movement initiation to develop. While this explanation is likely only part of the story, an exploratory analysis of our data confirms this impression, as we did indeed find that successful Stop-trials with fronto-central β-bursts had longer Stop-signal delays compared to trials without β-bursst (t(229) = 2.27, p = .024, d = .06)^1^.

In summary, we here provide the first report of β-burst activity underlying both movement initiation and cancellation in humans. Future studies should investigate the exact properties of these β-bursts as they relate to human and non-human behavior, movement disorders, and the neural basis of movement cancellation in cognitive control.

## Methods

### Participants

234 healthy adult humans (mean age: 22.7, SEM: .43, 137 female, 25 left-handed) from the Iowa City community participated in the study, either for course credit or for an hourly payment. 123 of those datasets were published as part of other studies, none of which focused on β-bursting [54, 61-63]. All procedures were approved by the local ethics committee at the University of Iowa (IRB #201511709).

### Task

The task was identical to the one described in [54, 61-63]. In short, trials began with a fixation cross (500ms duration), followed by a white left- or rightward arrow (Go-signal). Participants were instructed to respond as fast and accurately as possible to the arrow using their left or right index finger (the respective response-buttons were q and p on a QWERTY keyboard). On one-third of trials, a Stop-signal occurred (the arrow turned from white to red) at a delay after the go-stimulus (stop-signal delay, SSD). The SSD, which was initially set to 200ms, was dynamically adjusted in 50ms increments to achieve a p(stop) of .5: after successful stops, the SSD was prolonged; after failed stops, it was shortened. This was done independently for left- and rightward go-stimuli. Trial duration was fixed at 3,000ms. Six blocks of 50 trials were performed (200 Go, 100 Stop).

### Data availability

All data, procedures, and analysis routines can be downloaded on the Open Science Framework at [URL to be inserted after acceptance].

### Behavioral analysis

Means were extracted for each subject for the following measures: Go-trial reaction time, failed Stop-trial reaction time, Stop-signal delay, Stopping accuracy, and Stop-signal reaction time, which was calculated via the integration method [24].

### EEG recording

Scalp-EEG was recorded using two different BrainProducts (Garching, Germany) systems – one active (actiChamp) and one passive (MR plus). In both cases, 62-channel electrode caps with two additional electrodes on the left canthus (over the lateral part of the orbital bone of the left eye) and over the part of the orbital bone directly below the left eye were used. The ground was placed at electrode Fz, and the reference was placed at electrode Pz. EEG was digitized at a sampling rate of 500 Hz, with hardware filters set to 10s time-constant high-pass and 1000 Hz low-pass.

### EEG data preprocessing

Data were preprocessed using custom routines in MATLAB, incorporating functions from the EEGLAB toolbox [64]. The electrode * time-series matrices for each task were imported into MATLAB and then filtered using symmetric two-way least-squares finite impulse response filters (high-pass cutoff: .3 Hz, low-pass cutoff: 30 Hz). Non-stereotyped artifacts were automatically removed from further analysis using segment statistics applied to consecutive one-second segments of data [joint probability and joint kurtosis, with both cutoffs set to 5 SD, cf., 65]. After removal of non-stereotypic artifacts, the data were re-referenced to common average and subjected to a temporal infomax ICA decomposition algorithm [66], with extension to subgaussian sources [67]. The resulting component matrix was screened for components representing eye-movement and electrode artifacts using outlier statistics and non-dipolar components [residual variance cutoff at 15%, 68], which were removed from the data. The remaining components were subjected to further analyses. For all statistical analyses and all plots (except the topographical plots in Figure 1a and 2a), the data were subsequently re-referenced using the current-source density method [69], which minimizes the effects of volume conduction on the scalp-measured activity. This was done to permit β-event detection at fronto-central electrode FCz and lateral sensorimotor electrodes C3 and C4 without cross-contamination by either side.

### β-burst detection

β-burst detection was performed as described in Shin and colleagues’ work [3]. The description is adapted from therein.

First, each electrode’s data was convolved with a complex Morlet wavelet of the form:

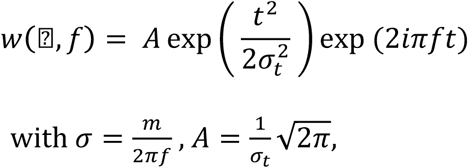

and *m* = 7 (cycles) for each of 15 evenly spaced frequencies spanning the β-band (15-29Hz). Time-frequency power estimates were extracted by calculating the squared magnitude of the complex wavelet-convolved data. These power estimates were then epoched relative to the events in question (ranging from -500 to +1,000ms with respect to Stop- / Go-signals). Individual β-bursts were defined as local maxima in the trial-by-trial β-band time-frequency power matrix for which the power exceeded a set cutoff of 6x the median power of the entire time-frequency power matrix for that electrode. Local maxima were identified using the MATLAB function imregional().

### Topographical distribution of β-bursts (Figures 1a, 2a, 3)

To visualize the topographical distribution of β-bursts with respect to Stop- and Go-signals, 12 windows of 25ms length, starting at 25ms after the event, were defined. For each subject, the number of β-bursts in each window at each electrode following the respective stimulus (Go/Stop) was counted. The average number of β-bursts in each time window for each of the three trial types (Correct Go, Successful Stop, Failed Stop) was then plotted in a topographical grid representing the scalp surface (Figures 1a and 2a). Figure 3 depicts the difference between the number of β-bursts on Go- and Stop-trials in each window and at each electrode, tested for significance using paired-samples t-tests. The resulting electrode * time window matrix of p-values was corrected for multiple comparisons to a significance level of p = .0001 using the false-discovery rate procedure (FDR, [70]).

### Temporal development of β-bursts (Figure 1b, 2b, 2d)

To visualize the temporal development of β-bursts on Go-trials at the two electrodes of interest (C3 and C4), 11 windows of 50ms length ranging from 25ms post-event to 575ms post-event were defined. This time range spanned the entire post-Go-signal period leading up to mean reaction time. To test the linear decreasing trend observed at C3/C4 during movement initiation for significance, we submitted the means for left- and right-hand responses to the Mann-Kendall test. To test the lateralization of the linear trend towards the end of the response period (i.e., prior to mean RT), we compared the means for left- and right-hand responses at both electrodes using paired-samples t-tests, again corrected for multiple comparisons to a significance level of p = .0001 using the false-discovery rate procedure. To visualize the temporal development of β-bursts on Stop-trials at the electrode of interest (FCz), 6 windows of 50ms length ranging from 25ms post-event to 325ms post-event were defined. This time range spanned the entire post-Stop-signal period leading up to Stop-signal reaction time.

### Comparison of pre-SSRT β-events (Figure 2c)

To test the difference in the amount of β-bursts at fronto-central electrode FCz for significance, we counted the number of β-bursts in the time period ranging from the Stop-signal to each individual participants’ SSRT estimate, separately for successful and failed Stop-trials. The means were compared across subjects using a paired-samples t-tests.

### Lateralized β-bursting after fronto-central β-bursts (Figure 4)

To investigate the pattern of β-bursting over bilateral electrodes C3/4 following fronto-central β-bursts in the Stop-Signal-to-SSRT period on successful Stop-trials, we identified each trial in which such a fronto-central β-burst event occurred, and counted the amount of β-bursts at electrodes C3/4 both contra- and ipsilateral to the to-be-stopped response in eight time windows of 25ms duration ranging from -100 to +100ms around the fronto-central β-burst event. In case more than one β-burst event was found, we chose the latency of the first of those events. To compare these β-burst counts to trials in which no fronto-central β-burst was found, a random time-point in the Stop-signal-to-SSRT interval was chosen from a uniform distribution and C3/4 β-bursts were quantified in an identical time window around that random time point in the pre-SSRT period. We then compared the means burst-counts for the two conditions in the four time-windows following the event (or the ‘pseudo-event’ in the case of the trial without an actual fronto-central burst) using signed-rank tests (the non-parametrical equivalent of the paired-samples t-test, chosen because of the large skew of the means towards zero in these samples), corrected for multiple comparisons to a critical p-value of p < .0001 using the FDR-method.

## Acknowledgements

The author would like to thank the following research assistants that were involved in collecting these data: Daniel Thayer, Julian Scheffer, Cailey Parker, Hailey Billings, Carly Ryder, Megan Hynd, Carly Iacullo, Brynne Dochterman, Isabella Dutra, Alec Mather, Kylie Dolan, Nathan Chalkley, Darcy Waller, Tobin Dykstra, and Cheol Soh. Furthermore, the author would like to thank Mark Blumberg, John Freeman, and Bob McMurray for helpful discussions. This research was supported by grants from the National Institute of Health (R01 NS102201), the National Science Foundation (NSF CAREER 1752355), and the Roy J Carver Foundation (Junior Research Program of Excellence).

## Footnotes

1 Note that the effect size is very small. This is likely an effect of the adaptive Stop-signal delay, which severely limits the variance in this analysis.

## References

1. Murthy, V.N. and Fetz, E.E. (1992) Coherent 25-to 35-Hz oscillations in the sensorimotor cortex of awake behaving monkeys. Proc Natl Acad Sci U S A 89 (12), 5670–4.

2. Sanes, J.N. and Donoghue, J.P. (1993) Oscillations in local field potentials of the primate motor cortex during voluntary movement. Proc Natl Acad Sci U S A 90 (10), 4470–4.

3. Shin, H. et al. (2017) The rate of transient beta frequency events predicts behavior across tasks and species.Elife 6.

4. Engel, A.K. and Fries, P. (2010) Beta-band oscillations--signalling the status quo? Curr Opin Neurobiol 20 (2), 156–65.

5. Pfurtscheller, G. et al. (2003) Spatiotemporal patterns of beta desynchronization and gamma synchronization in corticographic data during self-paced movement. Clin Neurophysiol 114 (7), 1226–36.

6. McFarland, D.J. et al. (2000) Mu and beta rhythm topographies during motor imagery and actual movements. Brain Topogr 12 (3), 177–86.

7. Swann, N. et al. (2011) Deep brain stimulation of the subthalamic nucleus alters the cortical profile of response inhibition in the beta frequency band: a scalp EEG study in Parkinson’s disease. J Neurosci 31 (15), 5721–9.

8. Swann, N. et al. (2009) Intracranial EEG reveals a time- and frequency-specific role for the right inferior frontal gyrus and primary motor cortex in stopping initiated responses. J Neurosci 29 (40), 12675–85.

9. Swann, N.C. et al. (2012) Roles for the pre-supplementary motor area and the right inferior frontal gyrus in stopping action: electrophysiological responses and functional and structural connectivity. Neuroimage 59 (3), 2860–70.

10. Wagner, J. et al. (2018) Establishing a Right Frontal Beta Signature for Stopping Action in Scalp EEG: Implications for Testing Inhibitory Control in Other Task Contexts. J Cogn Neurosci 30 (1), 107–118.

11. Picazio, S. et al. (2014) Prefrontal control over motor cortex cycles at beta frequency during movement inhibition. Curr Biol 24 (24), 2940–5.

12. Kuhn, A.A. et al. (2004) Event-related beta desynchronization in human subthalamic nucleus correlates with motor performance. Brain 127 (Pt 4), 735–46.

13. Ray, N.J. et al. (2012) The role of the subthalamic nucleus in response inhibition: evidence from local field potential recordings in the human subthalamic nucleus. Neuroimage 60 (1), 271–8.

14. Wessel, J.R. et al. (2016) Stop-related subthalamic beta activity indexes global motor suppression in Parkinson’s disease. Mov Disord 31 (12), 1846–1853.

15. Brittain, J.S. and Brown, P. (2014) Oscillations and the basal ganglia: motor control and beyond. Neuroimage 85 Pt 2, 637–47.

16. Hammond, C. et al. (2007) Pathological synchronization in Parkinson’s disease: networks, models and treatments. Trends Neurosci 30 (7), 357–64.

17. Jenkinson, N. and Brown, P. (2011) New insights into the relationship between dopamine, beta oscillations and motor function. Trends Neurosci 34 (12), 611–8.

18. Bronte-Stewart, H. et al. (2009) The STN beta-band profile in Parkinson’s disease is stationary and shows prolonged attenuation after deep brain stimulation. Exp Neurol 215 (1), 20–8.

19. Quinn, E.J. et al. (2015) Beta oscillations in freely moving Parkinson’s subjects are attenuated during deep brain stimulation. Mov Disord 30 (13), 1750–8.

20. Feingold, J. et al. (2015) Bursts of beta oscillation differentiate postperformance activity in the striatum and motor cortex of monkeys performing movement tasks. Proc Natl Acad Sci U S A 112 (44), 13687–92.

21. Leventhal, D.K. et al. (2012) Basal ganglia beta oscillations accompany cue utilization. Neuron 73 (3), 523–36.

22. Sherman, M.A. et al. (2016) Neural mechanisms of transient neocortical beta rhythms: Converging evidence from humans, computational modeling, monkeys, and mice. Proc Natl Acad Sci U S A 113 (33), E4885–94.

23. Logan, G.D. and Cowan, W.B. (1984) On the Ability to Inhibit Thought and Action: A Theory of an Act of Control. Psych Rev 91, 295–327.

24. Verbruggen, F. et al. (2019) A consensus guide to capturing the ability to inhibit actions and impulsive behaviors in the stop-signal task. Elife 8.

25. Boucher, L. et al. (2007) Inhibitory control in mind and brain: an interactive race model of countermanding saccades. Psychol Rev 114 (2), 376–97.

26. Verbruggen, F. and Logan, G.D. (2008) Response inhibition in the stop-signal paradigm. Trends Cogn Sci.

27. Rossiter, H.E. et al. (2014) Beta oscillations reflect changes in motor cortex inhibition in healthy ageing. Neuroimage 91, 360–5.

28. Doyle, L.M. et al. (2005) Lateralization of event-related beta desynchronization in the EEG during pre-cued reaction time tasks. Clin Neurophysiol 116 (8), 1879–88.

29. Kaiser, J. et al. (2001) Event-related beta desynchronization indicates timing of response selection in a delayed-response paradigm in humans. Neurosci Lett 312 (3), 149–52.

30. Kok, A. et al. (2004) ERP components associated with successful and unsuccessful stopping in a stop-signal task. Psychophysiology 41 (1), 9–20.

31. Wessel, J.R. and Aron, A.R. (2015) It’s not too late: the onset of the frontocentral P3 indexes successful response inhibition in the stop-signal paradigm. Psychophysiology 52 (4), 472–80.

32. Kenemans, J.L. (2015) Specific proactive and generic reactive inhibition. Neurosci Biobehav Rev 56, 115–26.

33. Swann, N.C. et al. (2011) Roles for the pre-supplementary motor area and the right inferior frontal gyrus in stopping action: Electrophysiological responses and functional and structural connectivity. Neuroimage.

34. Schmidt, R. et al. (2013) Canceling actions involves a race between basal ganglia pathways. Nature Neuroscience 16 (8), 1118–24.

35. Schall, J.D. et al. (2017) Models of inhibitory control. Philos Trans R Soc Lond B Biol Sci 372 (1718).

36. Badry, R. et al. (2009) Suppression of human cortico-motoneuronal excitability during the Stop-signal task. Clin Neurophysiol 120 (9), 1717–23.

37. Majid, D.S. et al. (2012) Transcranial magnetic stimulation reveals dissociable mechanisms for global versus selective corticomotor suppression underlying the stopping of action. Cerebral Cortex 22 (2), 363–71.

38. Wessel, J.R. et al. (2013) Saccade suppression exerts global effects on the motor system. Journal of Neurophysiology.

39. Duque, J. et al. (2017) Physiological Markers of Motor Inhibition during Human Behavior. Trends Neurosci 40 (4), 219–236.

40. Elchlepp, H. et al. (2016) Proactive inhibitory control: A general biasing account. Cogn Psychol 86, 27–61.

41. Greenhouse, I. et al. (2012) Stopping a response has global or nonglobal effects on the motor system depending on preparation. Journal of Neurophysiology 107 (1), 384–92.

42. Jaffard, M. et al. (2008) Proactive inhibitory control of movement assessed by event-related fMRI. Neuroimage 42 (3), 1196–206.

43. Stuphorn, V. and Emeric, E.E. (2012) Proactive and reactive control by the medial frontal cortex. Front Neuroeng 5, 9.

44. Verbruggen, F. and Logan, G.D. (2009) Proactive adjustments of response strategies in the stop-signal paradigm. J Exp Psychol Hum Percept Perform 35 (3), 835–54.

45. Vink, M. et al. (2015) The role of stop-signal probability and expectation in proactive inhibition. Eur J Neurosci 41 (8), 1086–94.

46. Anidi, C. et al. (2018) Neuromodulation targets pathological not physiological beta bursts during gait in Parkinson’s disease. Neurobiol Dis 120, 107–117.

47. Tinkhauser, G. et al. (2017) The modulatory effect of adaptive deep brain stimulation on beta bursts in Parkinson’s disease. Brain 140 (4), 1053–1067.

48. Gauggel, S. et al. (2004) Inhibition of ongoing responses in patients with Parkinson’s disease.J Neurol Neurosurg Psychiatry 75 (4), 539–44.

49. Obeso, I. et al. (2011) Deficits in inhibitory control and conflict resolution on cognitive and motor tasks in Parkinson’s disease. Exp Brain Res 212 (3), 371–84.

50. van den Wildenberg, W.P.M. et al. (2006) Stimulation of the subthalamic region facilitates the selection and inhibition of motor responses in Parkinson’s disease. Journal Cognitive Neuroscience.

51. Bartolo, R. and Merchant, H. (2015) beta oscillations are linked to the initiation of sensory-cued movement sequences and the internal guidance of regular tapping in the monkey. J Neurosci 35 (11), 4635–40.

52. Wessel, J.R. (2018) Surprise: A More Realistic Framework for Studying Action Stopping? Trends Cogn Sci 22 (9), 741–744.

53. Wessel, J.R. and Aron, A.R. (2017) On the Globality of Motor Suppression: Unexpected Events and Their Influence on Behavior and Cognition. Neuron 93 (2), 259–280.

54. Dutra, I. et al. (2018) Perceptual surprise improves action stopping by non-selectively suppressing motor activity via a neural mechanism for motor inhibition. J Neurosci.

55. Frank, M.J. et al. (2007) Hold your horses: impulsivity, deep brain stimulation, and medication in parkinsonism. Science 318 (5854), 1309–12.

56. Wessel, J.R. et al. (2019) Non-selective inhibition of inappropriate motor-tendencies during response-conflict by a fronto-subthalamic mechanism. Elife 8.

57. Wessel, J.R. (2018) An adaptive orienting theory of error processing. Psychophysiology 55 (3).

58. Ridderinkhof, K.R. (2002) Micro- and macro-adjustments of task set: activation and suppression in conflict tasks. Psychol Res 66 (4), 312–23.

59. Chikazoe, J. et al., Preparation to inhibit a response complements response inhibition during performance of a stop-signal task, J Neurosci, 2009, pp. 15870–7.

60. Bissett, P.G. and Logan, G.D. (2012) Post-stop-signal adjustments: inhibition improves subsequent inhibition. J Exp Psychol Learn Mem Cogn 38 (4), 955–66.

61. Wessel, J.R. (2017) Prepotent motor activity and inhibitory control demands in different variants of the go/no-go paradigm. Psychophysiology.

62. Wessel, J.R. (2016) A Neural Mechanism for Surprise-related Interruptions of Visuospatial Working Memory. Cereb Cortex.

63. Waller, D.A. et al. (2019) Common neural processes during action-stopping and infrequent stimulus detection: The frontocentral P3 as an index of generic motor inhibition. Int J Psychophysiol.

64. Delorme, A. and Makeig, S. (2004) EEGLAB: an open source toolbox for analysis of singletrial EEG dynamics including independent component analysis. J Neurosci Methods 134 (1), 9–21.

65. Delorme, A. et al. (2007) Enhanced detection of artifacts in EEG data using higher-order statistics and independent component analysis. Neuroimage 34 (4), 1443–9.

66. Bell, A.J. and Sejnowski, T.J. (1995) An information-maximization approach to blind separation and blind deconvolution. Neural Comput 7 (6), 1129–59.

67. Lee, T.W. et al. (1999) Independent component analysis using an extended infomax algorithm for mixed subgaussian and supergaussian sources. Neural Comput 11 (2), 417–41.

68. Delorme, A. et al. (2012) Independent EEG sources are dipolar. PLoS One 7 (2), e30135.

69. Tenke, C.E. and Kayser, J. (2005) Reference-free quantification of EEG spectra: combining current source density (CSD) and frequency principal components analysis (fPCA). Clin Neurophysiol 116 (12), 2826–46.

70. Benjamini, Y. et al. (2006) Adaptive linear step-up procedures that control the false discovery rate. Biometrika 93 (3), 491–507.

